# Fat body CLOCK restrains innate immunity and maintains survival under dietary stress in *Drosophila*

**DOI:** 10.64898/2026.07.20.739608

**Authors:** Keisuke Fukumura, Shorbon Mowla, Vinithra Kathirvel, Colin Fang, Shreya Shekar, Annika F. Barber

## Abstract

Both aging and a high-fat diet (HFD) dampen circadian gene transcription rhythms and promote chronic inflammation. How aging and HFD interact to affect peripheral molecular clocks and circadian behavior remains unclear. Using *Drosophila melanogaster*, we showed that aging and HFD additively dampened circadian behavior, with their adverse effects converging on the fat body (FB), a tissue that regulates systemic metabolism and innate immunity. Applying longitudinal *in vivo* bioluminescence recording in small, genetically defined cell populations, we found that molecular clocks in the FB were uniquely vulnerable to aging- and HFD-induced dampening of rhythm amplitude, whereas those in the clock neurons declined with age but were resistant to dietary stress. To test the consequences of this FB clock decline, we disrupted individual components of the core molecular clock specifically in the FB. We found that only CLOCK (CLK) disruption shortened lifespan on HFD, whereas disrupting its binding partner CYCLE (CYC), or the repressors PERIOD and TIMELESS, did not. Furthermore, CLK, but not CYC, disruption upregulated antimicrobial peptide expression in the FB, dampened behavioral rhythms, and suppressed locomotor activity, even though both CLK and CYC disruption comparably dampened clock gene oscillation in the FB. Together, these results indicate that FB CLK has a unique role in suppressing pro-inflammatory signals independently of CYC. Our findings provide insight into how stressors such as aging and HFD selectively disrupt the peripheral metabolic clock, and into the distinct roles of individual clock components, with implications for age-related inflammation and metabolic disease.

## INTRODUCTION

Age-associated pathologies such as metabolic dysfunction and chronic inflammation have become more prevalent as human lifespan has been substantially extended due to advances in public health and medicine^1,2^. A prominent feature of aging is the progressive decline of circadian rhythms, the approximately 24 h cycles in behavior and physiology that synchronize an animal’s internal state with environmental conditions such as light:dark cycles and food availability^3^. Circadian rhythms are tightly associated with both metabolism^4^ and immune function^5^. Chronic circadian disruption, as seen in shift workers, increases the risk of obesity and metabolic syndrome^6,7^, and circadian misalignment also elevates inflammatory markers^8^. In addition to aging, modern industrialized diets composed of a large amount of fat disrupt circadian rhythms^9^ and themselves induce chronic inflammation^10^.

At the molecular level, circadian rhythms are generated by a transcription-translation feedback loop (TTFL), a mechanism genetically and functionally conserved from insects to humans^11^. In *Drosophila*, the TTFL is composed of four core clock genes. The basic helix-loop-helix PAS-domain transcription factors CLOCK (CLK) and CYCLE (CYC) form a heterodimer and bind E-box elements to activate transcription of *period* (*per*) and *timeless* (*tim*), which encode transcriptional repressor proteins PER and TIM^12–14^. As PER and TIM accumulate, they form a complex and enter the nucleus to repress CLK:CYC transcriptional activity^14,15^. As PER and TIM are degraded, CLK:CYC resumes activating transcription, with this full cycle taking approximately 24 h. In the fly brain, approximately 240 clock neurons in which the TTFL operates, referred to as the central pacemaker, receive light input, coordinate behavioral rhythms, and synchronize molecular clocks in peripheral tissues^16,17^. The TTFL autonomously operates in most peripheral tissues to drive tissue-specific rhythmic gene transcription and function. While entrained by the central pacemaker, these peripheral clocks are also responsive to non-photic cues, including feeding, temperature, and stress signaling^18^.

Among insect peripheral tissues, the fat body (FB), which performs the functions of mammalian liver and adipose tissue, plays a principal role in metabolism and immune regulation^19^. Previous studies have shown that clock machinery in the FB is associated with nutrient storage and energy metabolism. FB-specific disruption of CLK increases food intake, decreases glycogen levels, and induces starvation intolerance^20^. Microarray analysis revealed that CLK disruption dampened the rhythmic expression of multiple genes involved in lipid, nucleotide, and carbohydrate metabolism, as well as detoxification, while time-restricted feeding drives rhythmic gene expression in the FB^21^. In addition to metabolic function, the FB is a primary site of humoral immune defense, producing antimicrobial peptides (AMPs) through the Toll and immune deficiency (IMD) pathways^22^. Although the regulatory mechanism of immune pathways remains to be characterized in the *Drosophila* FB, emerging evidence has shown that clock components regulate immune gene expression and immune response in a tissue-specific manner in mammals^23^, with the links to immune phenotypes observed in whole-body clock gene mutants in *Drosophila*^24,25^.

As in mammals, in *Drosophila*, metabolic stress is commonly induced by nutrient-excess diets, such as a high-fat diet (HFD). HFD increases body weight and lipid storage, alters immune gene expression, impairs locomotor performance, and shortens lifespan^26,27^. The detrimental effects of HFD vary with the source of dietary fat^27,28^. Similarly, aging alters systemic metabolism, increases AMP expression, and reduces locomotor performance^29–31^. Furthermore, aging and HFD are both known to impair the circadian clock. Aged *Drosophila* show weakened behavioral rhythms and fragmented sleep, as well as reduced amplitude of clock gene oscillation^32,33^. HFD has also been reported to dampen behavioral rhythms and alter sleep patterns^34,35^.

Despite this progress, the combined effects of aging and HFD on circadian behavior and molecular clocks, especially in peripheral metabolic tissues, and the consequences of peripheral clock disruption have not been characterized. We hypothesized that aging and HFD act together to dampen circadian behavior and molecular clocks in peripheral tissues, and that peripheral clocks are critical for maintaining lifespan and healthspan. To test this, we used *Drosophila melanogaster*, a model organism with a shorter lifespan and sophisticated genetic toolkits and found that aging and HFD additively dampened circadian behavior and converged on the FB clock, which was uniquely vulnerable to dietary stress compared to the central clock. Disruption of CLK in the FB, but not of other clock components, shortened lifespan on HFD, increased AMP expression, and impaired circadian behavior. These data reveal an additional function of CLK, separable from its timekeeping role, in restraining chronic immune activation. Our findings identify the FB clock as a principal site at which the detrimental effects of aging and HFD converge and establish CLK in the FB as a determinant of immune response and survival under dietary stress.

## RESULTS

### Age and high-fat diet individually dampen locomotor activity and rhythms, and interact to affect sleep architecture

Because both aging and HFD disrupt circadian behavioral rhythms, we sought to characterize their interactions by aging flies on a HFD. We first analyzed circadian behavior and sleep architecture in young (7–9 days old) and middle-aged (21–23 days old) flies on a standard diet (SD) or a lard-based HFD using the Drosophila Activity Monitor (DAM) system. Visualization of the averaged circadian activity profile in constant darkness (DD) clearly demonstrates that aging and diet each reduce the amplitude of daily activity peaks (Fig. 1A). We used two-way ANOVA to analyze the effects of age, diet, and their interaction on circadian behavioral parameters and found that both aging and exposure to HFD individually decreased behavioral rhythm strength assessed by fast Fourier transform (FFT) and 24 h activity with no age × diet interaction (age *P* < 0.0001, diet *P* < 0.0001, interaction n.s.; Fig. 1B, C). Neither age nor diet affected period length (Fig. 1D). Visualization of the average sleep profile in a 12 h light :12 h dark (LD) cycle shows only small effects of aging and HFD on sleep amount (Fig. 1E). However, the interaction of aging and HFD showed significant effects on sleep architecture (Fig. 1 F–G). Two-way ANOVA showed a modest effect of diet on 24 h sleep amount, with HFD decreasing sleep only in young flies (age n.s., diet *P* < 0.01, interaction *P* < 0.01 Fig. 1F). In contrast, both aging and HFD increased the number of sleep bouts (age *P* < 0.0001, diet *P* < 0.01, interaction *P* < 0.01; Fig. 1G) and decreased sleep bout duration (age *P* < 0.0001, diet *P* < 0.001, interaction *P* < 0.001; Fig. 1H), indicating fragmented sleep. Consistent with the age × diet interactions in the effect on sleep bout number and duration, HFD did not further exacerbate sleep fragmentation in middle-aged flies (Fig. 1G, H), which already showed fragmented sleep. These data demonstrate that aging and HFD additively disrupt circadian rest-activity rhythms while interacting to affect sleep architecture.

**Fig. 1.**
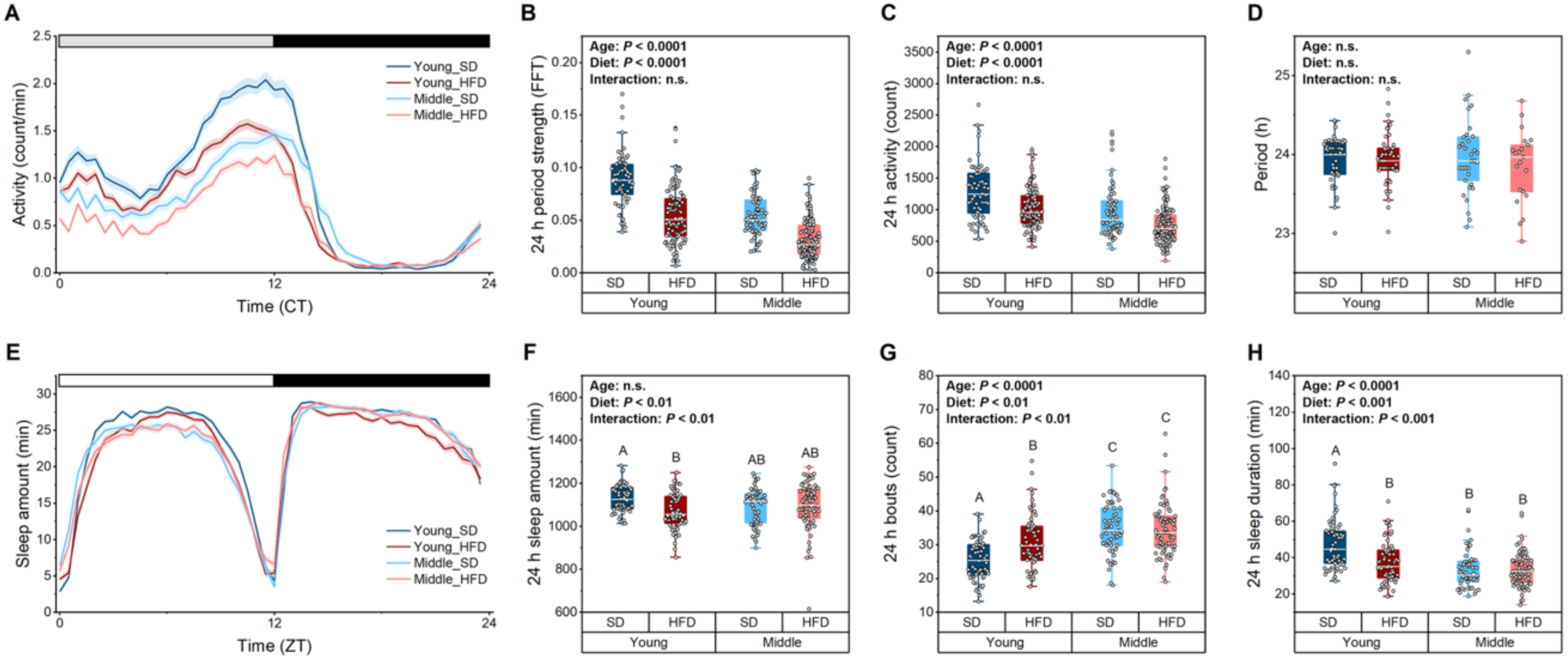
Age and high-fat diet interact to affect behavioral rhythms and sleep architecture. (A) Average locomotor activity per minute across 24 h from days 2–7 of constant darkness (DD) for young (7–9 days) and middle-aged (21–23 days) male flies maintained on a standard diet (SD) or a high-fat diet (HFD): young SD (dark blue), young HFD (dark red), middle-aged SD (light blue), and middle-aged HFD (light red). Lines and shaded areas represent the mean ± SEM. The horizontal bar denotes the lighting conditions during subjective day (open) and subjective night (filled) under DD. (B–D) Circadian behavior parameters: 24 h period strength by FFT (B), total 24 h activity counts (C), and free-running period by chi-squared periodogram (D) calculated from day 2–7 in DD. (E) Average sleep time per 30 minutes across 24 h from days 1–6 of a 12:12 light-dark cycle (LD). Lines and shaded areas represent the mean ± SEM respectively. The horizontal bar denotes the light (open) and dark (filled) phases under LD. (F–H) Sleep architecture parameters: Total 24 h sleep amount (F), number of sleep bouts per 24 h (G), and average sleep bout duration (H). For all box plots, the box spans the interquartile range (25th–75th percentiles), the central line is the median, whiskers extend to 1.5 times the interquartile range, and each point is an individual fly. Groups that do not share a common letter differ significantly (*P* < 0.05) by two-way ANOVA with age and diet as factors followed by Tukey’s *post hoc* test. *P* values for each main effect and their interaction are shown in each panel (n.s., not significant). Groups were compared by Tukey’s *post hoc* test when the interaction was significant (F–H). n = 63–121 (A–C), 21–59 (D), and 56–73 (E–H) flies per group across two independent replicates.

### FB clock gene oscillation is uniquely sensitive to diet-induced stress

To determine how aging and HFD affect the circadian system at the molecular level, we assessed rhythmic transcription of the core clock genes *Clk*, *per*, and *tim* by qRT-PCR in the head and FB across circadian time (CT). On SD, young and middle-aged flies exhibited robust oscillations of C*lk*, *per*, *tim* in the head (JTK-Cycle *P* < 0.05; Fig. 2A–C). In contrast, in the FB the amplitude of these clock transcript rhythms significantly dampened with age (*clk*, 1.775 to 0.179; *per*, 1.006 to 0.158; *tim*, 0.889 to 0.260), to the point that *per* transcription became arrhythmic in middle-aged flies (Fig. 2D–F). Similarly, rhythmic transcription was maintained in the heads of young flies on both SD and HFD (JTK_Cycle *P* < 0.05; Fig. 2G–I), whereas in the FB the amplitude was reduced on HFD compared to SD (*clk*, 1.775 to 0.507; *per*, 1.006 to 0.290; *tim*, 0.889 to 0.439), with *per* transcription again becoming arrhythmic (Fig. 2J–L). Although *Clk* expression remains rhythmic on HFD, albeit at reduced amplitude, the timing of peak expression is shifted to CT 4 on HFD compared to CT 12 on SD (Fig. 2J). We did not measure *cyc* because, unlike other clock genes, *cyc* is constitutively expressed and show no rhythmicity at the mRNA level^13^.

**Fig. 2.**
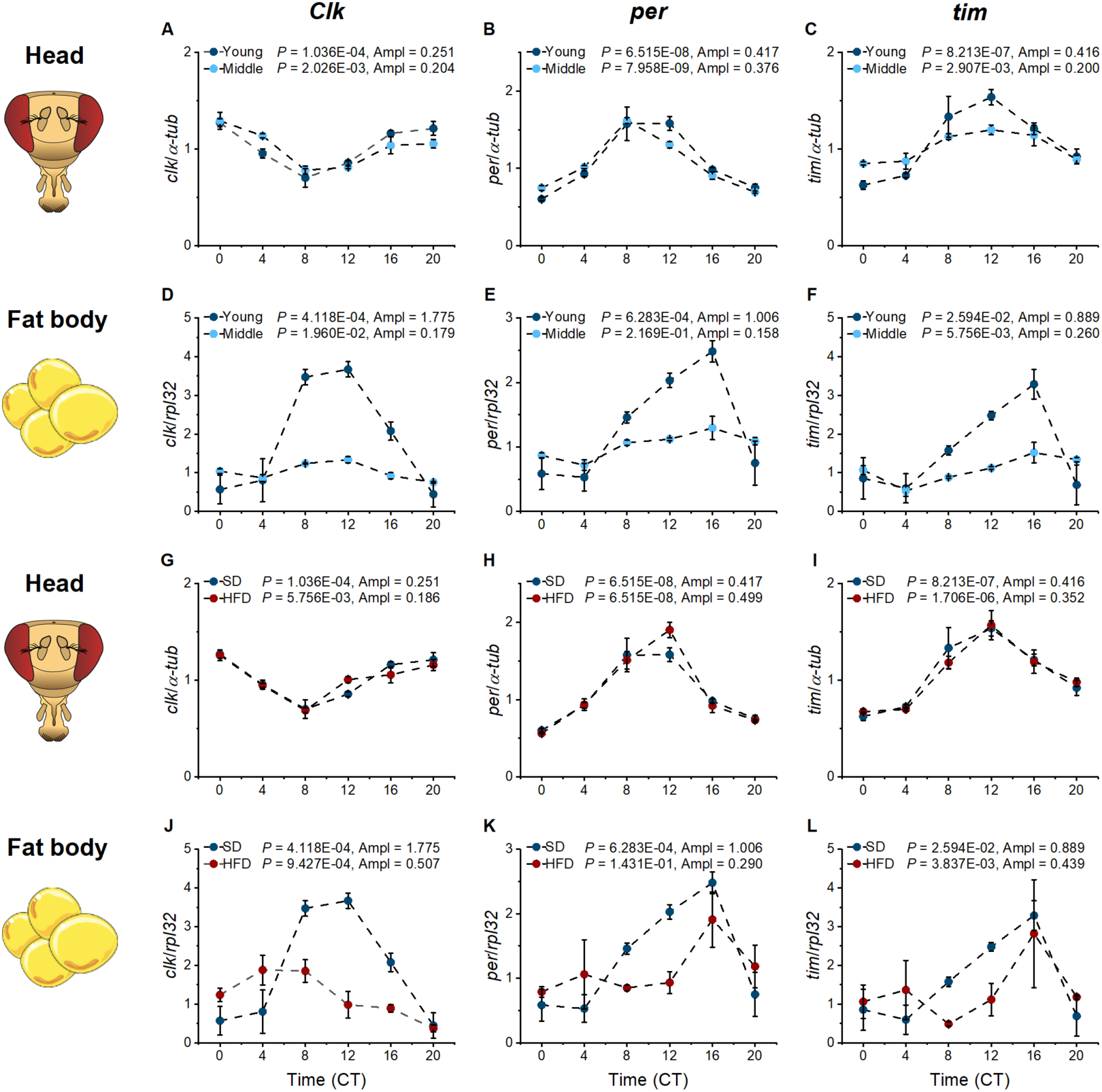
Aging or high-fat diet dampen the rhythmic transcription of core clock genes in the fat body. Daily oscillation of clock gene expression in the head (A–C) and fat body (FB; D–F) of young (7–9 days old; dark blue) and middle-aged (28–30 days old; light blue) male flies maintained in constant conditions on a standard diet (SD) determined by qRT-PCR. mRNA expression rhythms of *Clock* (*Clk*; A, D), *period* (*per*; B, E), and *timeless* (*tim*; C, F) were unaffected by age in heads while in fat body rhythm amplitude dampened with age. Daily oscillation of clock gene transcripts in the head (G–I) and FB (J–L) of young male flies maintained in constant conditions on SD (dark blue) or a high-fat diet (HFD; dark red). mRNA expression rhythms of *Clk* (G, J), *per* (H, K), and *tim* (I, L) were unaffected by HFD in heads while in fat body rhythm amplitude dampened on HFD. Expression in the head and FB was normalized to *α-tubulin* (*α-tub*) and *ribosomal protein L32* (*rpl32*), respectively. Data represent the mean ± SEM of 3 biological replicates per time point; each head replicate comprised 50 heads, and each FB replicate comprised 15 FBs. Rhythmicity (*P*) and amplitude (Ampl) were determined by JTK_Cycle.

To further interrogate the combined effects of aging and HFD on the circadian TTFL, we implemented the LABL (Locally Activatable BioLuminescence) reporter as a longitudinal approach with high resolution in defined tissues. Expression of tissue-specific FLP recombinase places luciferase under control of the *per* promoter (*per*-luc) so that bioluminescence reflects *per* transcription in real time in living flies^36^. We drove LABL expression in the FB using *Lsp2*-Gal4 and in a subset of clock neurons using *dvPDF*-Gal4 which is expressed in the ventrolateral (LNv) and dorsal lateral (LNd) neurons. Flies were reared on SD and randomly assigned to SD or HFD at eclosion. Bioluminescence was recorded across three age windows (7–16, 14–23, and 21–30 days post-eclosion). Based on previous studies showing that neurons are less sensitive to changes in nutritional state^37,38^, we hypothesized that the FB clock would be highly sensitive to diet-induced effects, whereas the brain clock neurons would be comparatively resistant. In the FB, we found that aging on SD reduced the amplitude of *per*-luc oscillation over the course of 7–30 days without affecting period length (Fig. 3A–D, H). Exposure to HFD reduced *per*-luc oscillation compared to SD at the youngest (7–16 days) window (Fig. 3A, D, E). Aging on HFD did not induce further decline in *per*-luc oscillation (Fig. 3D–G), which we hypothesize is due to a floor effect as the *per*-luc amplitude is already dampened at the youngest window. Two-way ANOVA supported this hypothesis, revealing a significant effect of diet and a significant age × diet interaction on *per*-luc amplitude (age n.s., diet *P* < 0.006, interaction *P* < 0.001; Fig. 3D). Period length was unaffected by HFD over the course of 7–30 days (Fig. 3H). In the clock neurons, the amplitude of *per*-luc oscillation dampened with aging with no effect on period length (Fig. 3I–L, P) as previously reported^36^. As in fat body, HFD reduced the amplitude of *per*-luc oscillation compared to SD within the 7–16 day window (Fig 3 L, M). However, we did not find any significant difference between SD and HFD within 14–23 and 21–30 day windows (Fig. 3L–O). Two-way ANOVA also indicated a significant effect of age, but not diet, and a significant age × diet interaction (age *P* < 0.002, diet n.s., interaction *P* < 0.0009; Fig. 3L). Comparing the amplitude differences between SD and HFD in the FB (Fig. 3D) and the clock neurons (Fig. 3L) within the 7–16 day window showed a larger diet-induced amplitude dampening in the FB compared to the clock neurons. Thus, we conclude that the TTFL in the clock neurons is sensitive to aging but resistant to HFD, while the TTFL in the FB is vulnerable to both aging and HFD.

**Fig. 3.**
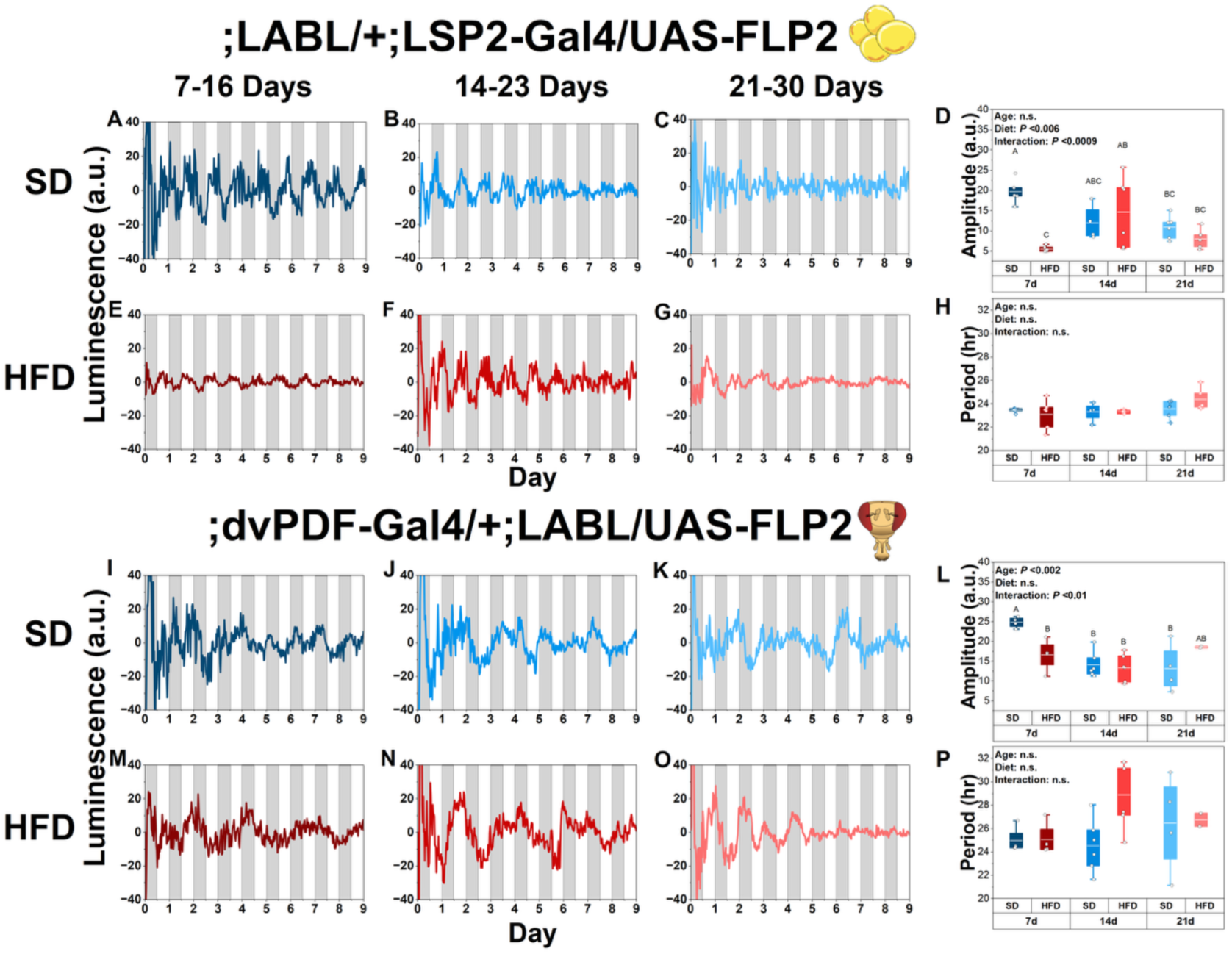
Fat body *period* gene transcription is more sensitive to age- and diet-induced dampening than brain clock neurons. (A–C) *In vivo period-*luciferase (*per-*luc) oscillations driven by *Lsp2-*Gal4 in the fat body over 9 days on a standard diet (SD) for flies loaded at 7–16 days (A), 14–23 days (B), and 21–30 days post-eclosion (C). (D) Quantification of fat body *per-*luc rhythm amplitude for flies on SD (blue) and a high-fat diet (HFD; red). HFD reduced *per-*luc rhythm amplitude at the youngest window and produced no further decline with age. (E–G) As in (A–C) for flies maintained on HFD from day 3 post-eclosion and fat body *per-*luc recorded from 7–16 days (E), 14–23 days (F), and 21–30 days. Quantification of fat body *per-*luc period length for flies on SD (blue) and HFD (red). (I–K) As in (A–C) for *per-*luc oscillations driven by *dvPDF-*Gal4 in brain clock neurons over 9 days on SD for flies loaded at 7–16 days (I), 14–23 days (J), and 21–30 days post-eclosion (K). (L) Quantification of brain clock *per-*luc rhythm amplitude for flies on SD (blue) and HFD (red). The amplitude of *per-*luc oscillation dampened with age. (M–O) As in (A–C) for flies maintained on HFD from day 3 post-eclosion and brain clock *per-*luc recorded from 7–16 days (M), 14–23 days (N), and 21–30 days post-eclosion (O). (P) Quantification of brain clock *per-*luc period length for flies on SD (blue) and HFD (red). For (A– C), (E–G), (I–K), and (M–O), gray and white vertical areas denote subjective night and subjective day, respectively. For all box plots, the box spans the interquartile range (25th–75th percentiles), the central line is the median, whiskers extend to 1.5 times the interquartile range, and each point represent a single recording (a chamber with 15 flies). Groups that do not share a common letter differ significantly (*P* < 0.05) by two-way ANOVA with age and diet as factors followed by Tukey’s *post hoc* test. *P* values for each main effect and their interaction are shown in each panel (n.s., not significant). Groups were compared by Tukey’s *post hoc* test when the interaction was significant (F–H). n = 5–6 (D and H) or 2–6 (L and P) chambers per group across at least two independent replicates.

### FB CLK, but not CYC, disruption impairs survival on HFD

To determine the physiological consequences of peripheral circadian TTFL dysfunction, we disrupted core clock genes specifically in the FB using *Lsp2*-Gal4 driver to overexpress a dominant-negative form of CLK (*Lsp2-*Gal4>*UAS*-CLK^DN^) or CYC (*Lsp2-*Gal4>*UAS*-CYC^DN^) protein, or to induce CRISPR-mediated knockout of *per* (*Lsp2-Gal4>UAS-per*^CRISPR^) or *tim* (*Lsp2-Gal4>UAS-tim*^CRISPR^), and measured survival rate on SD or HFD. On SD, none of these disruptions induced any significant lifespan changes compared to both parental genetic controls (Fig. 4A–D). In contrast, on HFD, only *Lsp2-*Gal4>*UAS*-CLK^DN^ markedly decreased survival compared to both parental controls, whereas *Lsp2-*Gal4>*UAS*-CYC^DN^, *Lsp2-*Gal4*>UAS-per*^CRISPR^, and *Lsp2-*Gal4*>UAS-tim*^CRISPR^ increased it (Fig. 4E–H). Given that CLK and CYC dimerize to serve as a transcriptional activator^11^, the opposing effects of CLK and CYC disruption prompted us to confirm whether the manipulations were effective. We found that both *Lsp2-*Gal4>*UAS*-CLK^DN^ and *Lsp2-*Gal4>*UAS*-CYC^DN^ dampened the amplitude of clock gene oscillation in the fat body in constant conditions (Fig. S1A–C), confirming that the TTFL is impaired in these flies. These data imply that CLK has a role in regulating HFD tolerance separable from its CYC-dependent role in the circadian TTFL.

**Fig. 4.**
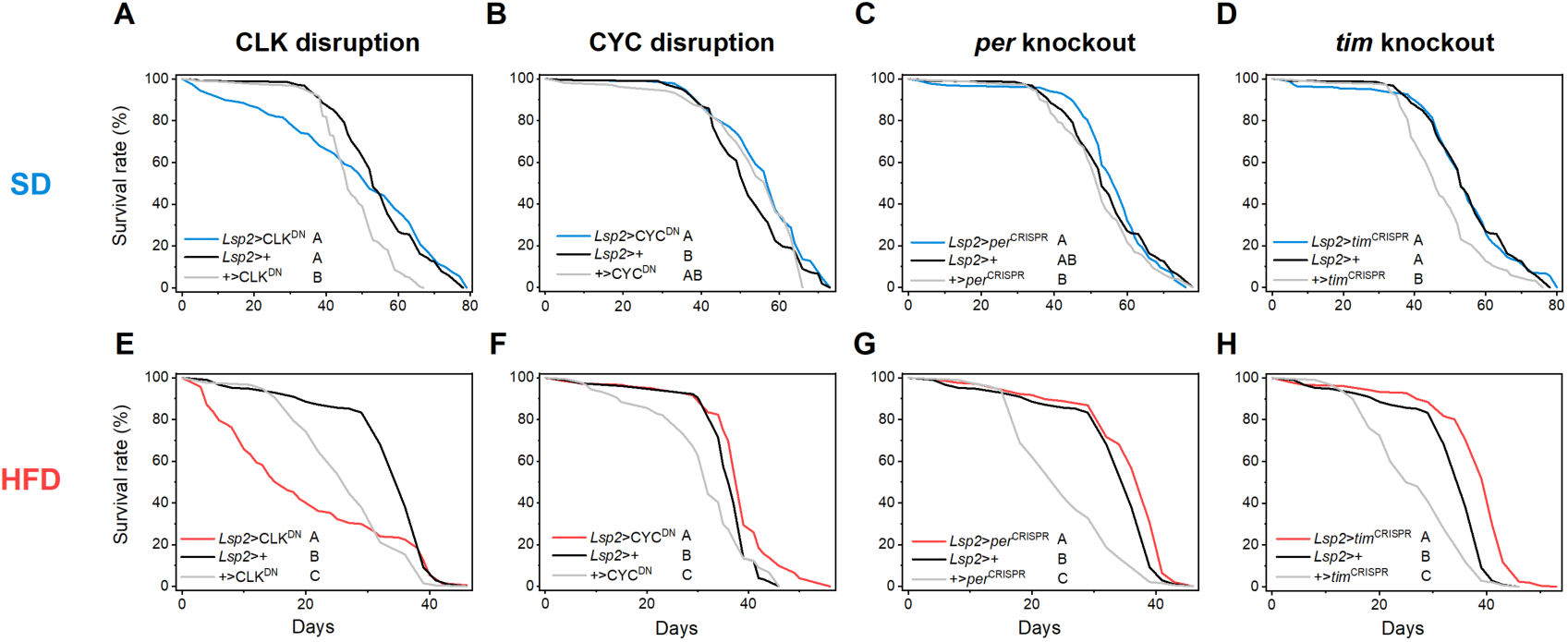
Fat body CLOCK disruption reduces lifespan on high-fat diet. Kaplan-Meier survival curves of flies with fat body-specific disruption of clock genes on a standard diet (SD; A–D) or a high-fat diet (HFD; E–H). Disruption of CLK (CLK^DN^), CYC (CYC^DN^), *per* (*per*^CRISPR^), or *tim* (*tim*^CRISPR^) with *Lsp2*-*Gal4* had no significant effect on survival on SD relative to both genetic controls (A–D), whereas CLK^DN^ reduced survival rate on HFD (E). In contrast, CYC^DN^, *per*^CRISPR^, and *tim*^CRISPR^ increased survival rate on HFD (F–H). In each panel, the experimental genotype is compared with the Gal4 driver control (*Lsp2*>+) and the corresponding UAS parental control (+>transgene). Groups that do not share a common letter differ significantly (*P* < 0.05) by pairwise log-rank test. *n* = 195–225 flies per group across two independent replicates.

### FB CLK disruption induces overactivation of immune pathways

The differences in lifespan between CLK vs. CYC disruption in the fat body were unexpected, so we sought to identify downstream gene expression differences that may underlie the differences in lifespan on HFD. To do this, we performed RNA-seq on the FBs of young (7-9 day) male *Lsp2-*Gal4>*UAS*-CLK^DN^ and *Lsp2-*Gal4>*UAS*-CYC^DN^ flies maintained on HFD and compared gene expression to *Lsp2-Gal4*>+ parental genetic controls. We found that CLK disruption induced large transcriptional changes with genes both up- and down-regulated (Fig. 5A). With stringent fold-change and expression abundance cutoffs (|log_2_ fold change| > 2.5, mean expression A > 6), we obtained 781 up- and 32 down-regulated candidate genes in *Lsp2-*Gal4>*UAS*-CLK^DN^, whereas *Lsp2-*Gal4>*UAS*-CYC^DN^ yielded only a few (5 up- and 1 down-regulated genes), with no overlap between the two manipulations (Fig. 5B). Functional enrichment analysis using Gene Ontology (GO) revealed an immune-related signature in both up- and down-regulated genes in *Lsp2-*Gal4>*UAS*-CLK^DN^, with the up-regulated candidates enriched for GO:0019731: antibacterial humoral response and the down-regulated candidates for GO:0002376: immune system process (Fig. 5C, D). Up-regulated genes also included genes with GO terms associated with cilium assembly and spermatid development (Fig. 5C). These genes likely represent contamination from male reproductive tissues in the manual FB dissection process. Furthermore, we found that CLK, but not CYC, disruption elevated transcripts of the antimicrobial peptides *Defencin, Drosocin, Diptericin A, Drosomycin,* and *Listericin*, whereas it reduced *Turandot A,* a gene that protects against immune overactivation (Fig. 5E–J). Additional immune genes were also upregulated when CLK was disrupted, but did not reach the same level of significance. The same immune-related signature was observed in an independent RNA-seq experiment on SD (Fig. S2A–F). Thus, we conclude that CLK, but not CYC, has a specific function in restraining immune-related processes on SD and HFD. Consequently, loss of CLK function in the FB drives a pro-inflammatory gene expression state that may contribute to the reduced lifespan in the presence of the additional inflammatory environmental stressor of HFD.

**Fig. 5.**
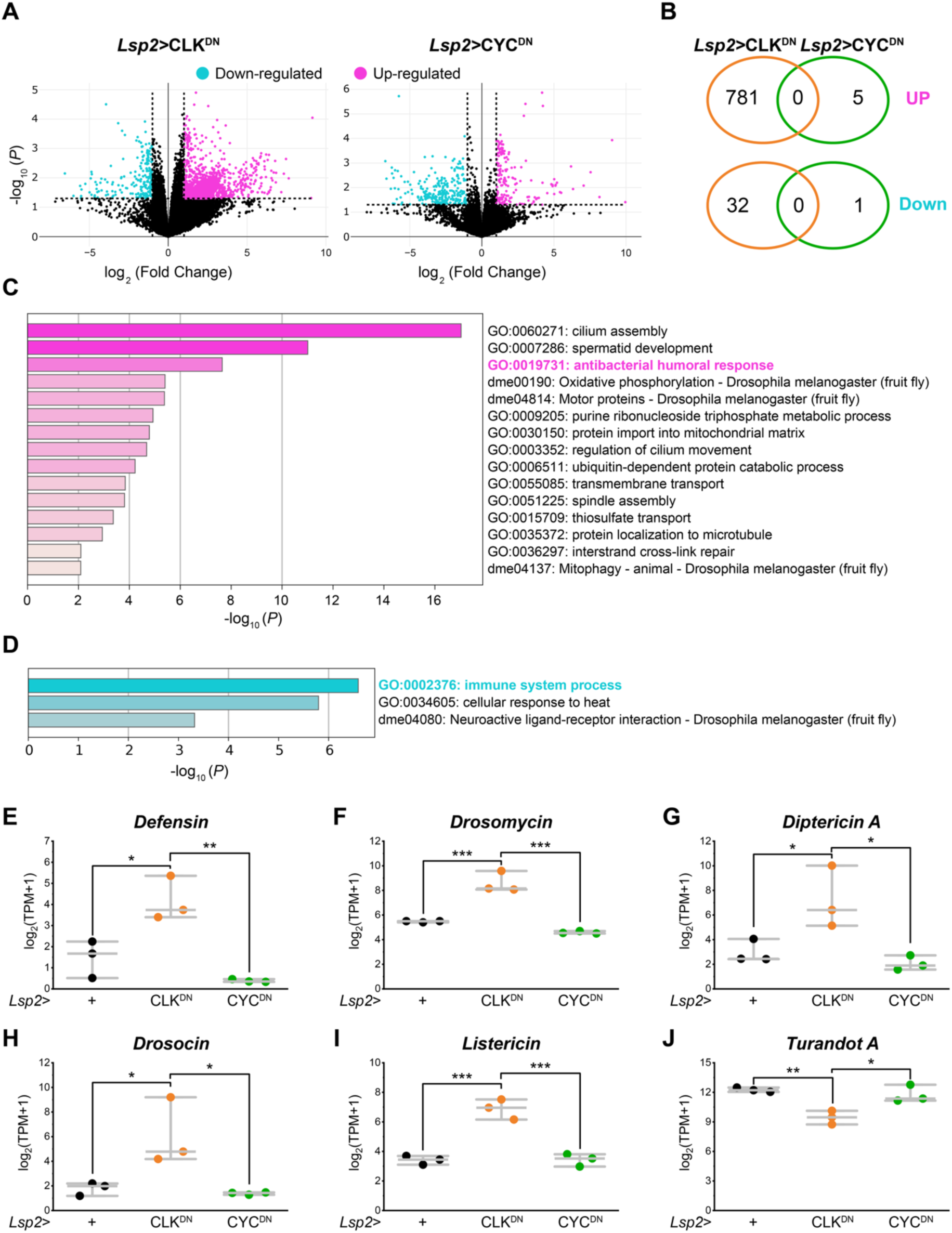
Disruption of CLOCK, but not CYCLE, induces overactivation of immune pathways. (A) Volcano plots of differential gene expression in the fat bodies of young (7–9 days old) male flies maintained on a high-fat diet (HFD): *Lsp2-*Gal4>*UAS*-CLK^DN^ (left) and *Lsp2-*Gal4>*UAS*-CYC^DN^ (right) flies, compared to the *Lsp2-*Gal4>+ control. The x- and y-axes show log_2_ fold change and −log_10_ *P* value, respectively. Magenta and cyan dots are significantly up- and down-regulated genes respectively (|log2 fold change| > 1 and *P* < 0.05; dashed lines), and black points are not significant. (B) Venn diagrams of the up- (top) and down-regulated (bottom) genes in *Lsp2-*Gal4>*UAS*-CLK^DN^ (orange) and *Lsp2-*Gal4>*UAS*-CYC^DN^ (green); the numbers of unique and shared genes are defined with more stringent cutoff (|log2 fold change| > 2.5 and mean expression A > 6). (C, D) Metascape functional enrichment of the genes up- (C) and down-regulated (D) in *Lsp2-*Gal4>*UAS*-CLK^DN^, ranked −log_10_ *P*; Gene Ontology terms (GO:) and KEGG pathways (dme) are shown, with immune-related terms highlighted. (E–J) Expression of antimicrobial peptide and immune genes: *Defensin* (E), *Drosomycin* (F), *Diptericin* A (G), *Drosocin* (H), *Listericin* (I), and *Turandot A* (J), in *Lsp2-*Gal4 controls (black), *Lsp2-*Gal4>*UAS*-CLK^DN^ (orange), and *Lsp2-*Gal4>*UAS*-CYC^DN^ flies (green). All immune genes were elevated in *Lsp2-*Gal4>*UAS*-CLK^DN^, except for *Turandot A* (J), which was reduced, whereas *Lsp2-*Gal4>*UAS*-CYC^DN^ had no significant effect. In (E–J), top and bottom horizontal lines indicate interquartile range (25th–75th percentiles), the central line is the median, and each point is one biological replicate. *n* = 3 per genotype; each FB replicate comprised 20 FBs. Data in (E–J) were analyzed by one-way ANOVA followed by Tukey’s *post hoc* test. *, *P* < 0.05; **, *P* < 0.01; ***, *P* < 0.001.

### FB CLK disruption dampens locomotor activity and rhythms without affecting sleep

Because systemic activation of the immune system is associated with behavioral declines across species^39,40^, we further asked whether CLK disruption in the FB induces behavioral changes. Using the DAM assay, we found that CLK disruption significantly decreased locomotor activity (Fig. 6A, C) and dampened behavioral rhythms (Fig. 6B) in young (7–9 day) male flies on SD, whereas CYC disruption did not affect these behavioral phenotypes. Period length was unaffected in both *Lsp2-*Gal4>*UAS*-CLK^DN^ and *Lsp2-*Gal4>*UAS*-CYC^DN^ flies (Fig. 6D). Unlike our findings with aging and dietary stress in our control line (Fig 1 E–H), sleep architecture was largely preserved. *Lsp2-*Gal4>*UAS*-CLK^DN^ and *Lsp2-*Gal4>*UAS*-CYC^DN^ flies showed no significant changes in total sleep amount (Fig. 6E, F), the number of sleep bouts (Fig. 6G), or sleep bout duration (Fig. 6H). These findings indicate that CLK, but not CYC, disruption impairs circadian locomotor activity rhythms without affecting sleep, mirroring CLK-specific immune upregulation even in the absence of dietary stress.

**Fig. 6.**
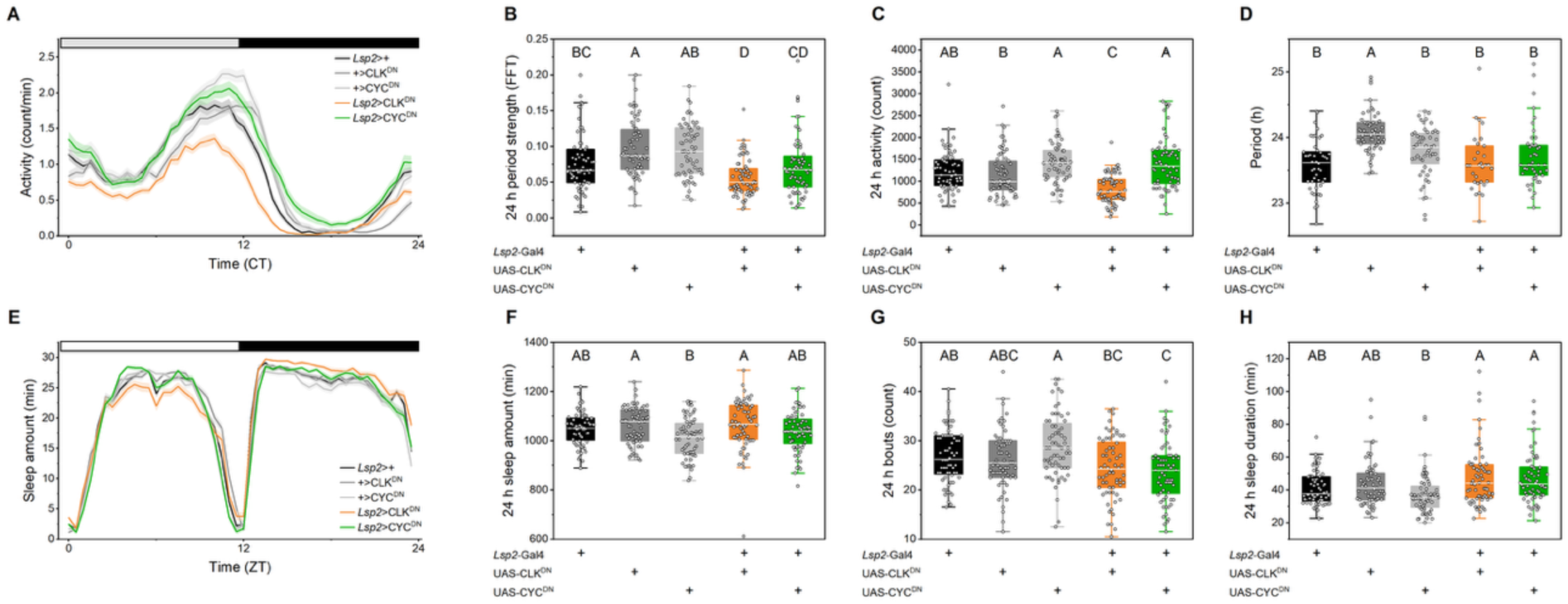
Fat body CLOCK disruption dampens behavioral rhythms and activity. (A) Average locomotor activity profile per minute across 24 h from days 2–7 of constant darkness (DD) for young (7–9 days old) male flies maintained on a standard diet for Gal4 control (*Lsp2*-Gal*4*>+; black), UAS controls (+>*UAS*-CLK^DN^; dark gray, +> *UAS*-CYC^DN^; light gray), and *Lsp2*-Gal4>*UAS*-CLK^DN^ (orange) and *Lsp2*-Gal4>*UAS*-CYC^DN^ (green). Lines and shaded areas represent the mean ± SEM. The horizontal bar denotes the subjective day (open) and subjective night (filled) under constant darkness. (B–D) Circadian behavior parameters: 24 h period strength by FFT (B), total 24 h activity (C), and free-running period by chi-squared periodogram (D) calculated from day 2–7 in DD. (E) Average sleep time per 30 minutes across 24 h from days 1–3 of a 12:12 light–dark cycle (LD). Lines and shaded areas represent the mean ± SEM. The horizontal bar denotes the light (open) and dark (filled) phases under LD. (F–H) Sleep architecture parameters: total 24 h sleep amount (F), number of sleep bouts per 24 h (G), and average sleep bout duration (H). For all box plots, the box spans the interquartile range (25th–75th percentiles), the central line is the median, whiskers extend to 1.5 times the interquartile range, and each point is an individual fly. Groups that do not share a common letter differ significantly (*P* < 0.05) by two-way ANOVA with age and diet as factors followed by Tukey’s *post hoc* test. n = 60–64 (A–D), 31–58, and 63–64 (E–H) flies per group across 2 independent replicates.

## DISCUSSION

Aging and HFD both dampen circadian rhythms and disrupt molecular clock function, yet whether and how they interact to affect rhythms and the molecular clock, especially in peripheral metabolic tissues, remains to be understood. In our study using *Drosophila melanogaste*r, we found that aging and HFD, individually, dampened behavioral rhythms and reduced locomotor activity, while interacting to affect sleep architecture (Fig. 1). Furthermore, at molecular level, we identified the *Drosophila* FB clock as being uniquely vulnerable to aging and HFD (Figs. 2, 3). Disruption of CLK in the FB elevated immune gene expression, reduced survival on HFD but not SD, and impaired behavioral rhythms and activity without affecting sleep architecture (Figs. 4, 5, 6). Importantly, the other clock gene disruptions did not reproduce the effects of CLK disruption, even though they too disrupted molecular clock oscillation, implying additional functions of CLK separable from its canonical role in the TTFL. Our findings suggest a model (Fig. 7) in which aging and HFD converge on the FB clock, and in which CLK regulates immune and metabolic homeostasis.

**Fig. 7.**
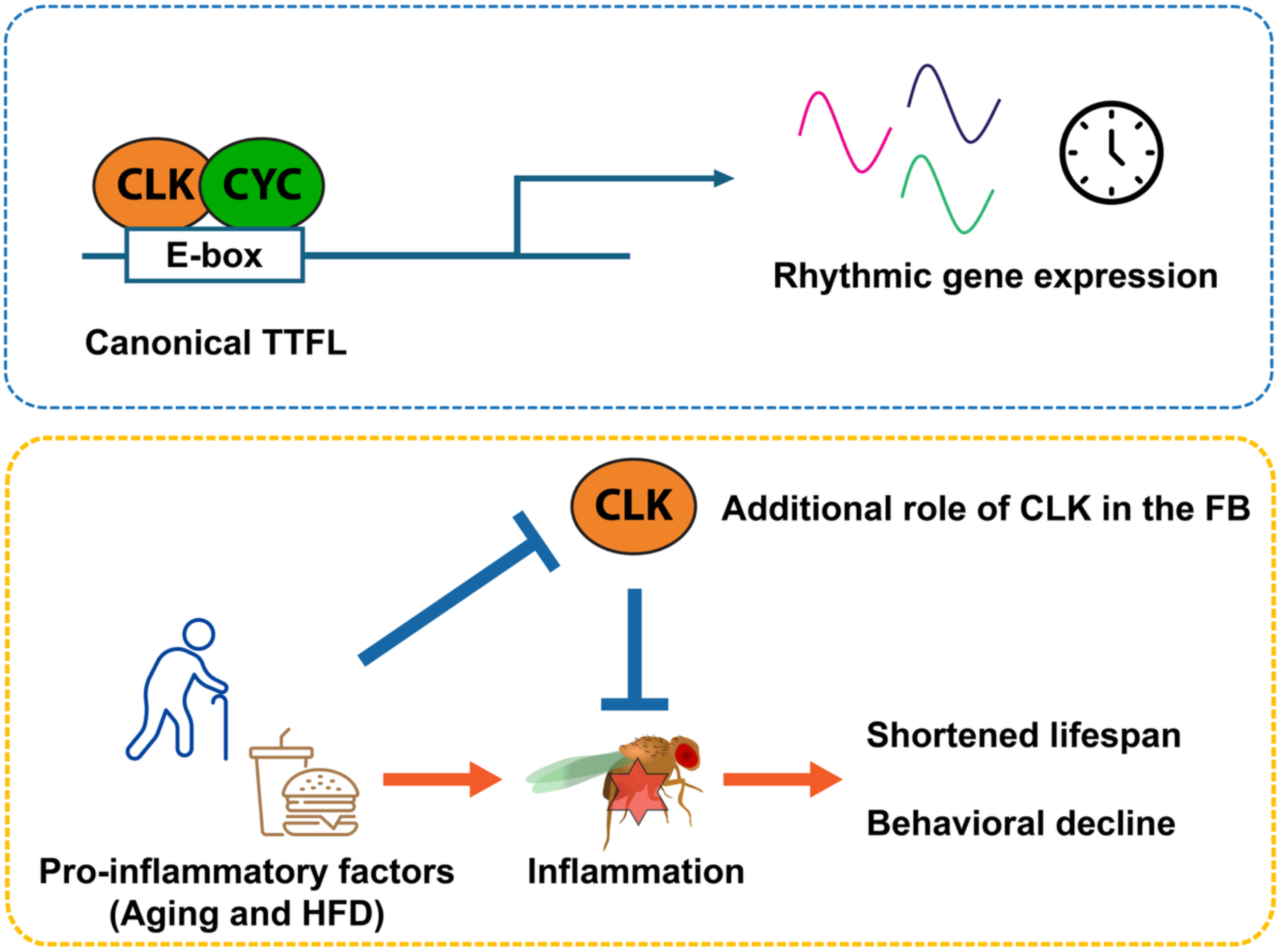
Proposed model. In the transcription-translation feedback loop (TTFL), CLK and CYC act together to bind E-box elements and activate rhythmic gene expression. We propose a model in which pro-inflammatory factors including aging and high-fat diet (HFD) induce inflammation in the fat body (FB), shortening lifespan and causing behavioral decline. Separable from its timekeeping role in the canonical TTFL, CLK also acts in the FB to restrain the inflammation and preserve survival and behavior.

Among the tissues we tested, the FB clock was uniquely vulnerable to stressors such as aging and HFD. By qRT-PCR, we found that the amplitude of clock gene oscillation in the FB was dampened by both aging and HFD, whereas the oscillation in the head remained robust (Fig. 2). Longitudinal measurement of *per* oscillation with the LABL assay^36^, which can resolve small cell populations, showed that HFD rapidly dampened *per* oscillation in the FB more than in the LNv and LNd clock neurons, identifying the FB clock, rather than the central pacemaker, as a principal target of dietary stress (Fig. 3). The LABL assay also showed that *per* oscillation in these clock neurons was dampened with age. The robustness of the whole-head clock genes measured by qRT-PCR and this age-related neuronal decline of *per* appear contradictory but are reconciled once the cellular makeup of the head is considered. The brain contains only ∼240 clock neurons that express the core clock TTFL^17^, greatly outnumbered by other head tissues such as the glia and visual tissue, so their decline is likely masked in whole-head measurements. These data further indicate that the clock gene oscillation declines earlier in the clock neurons than in the bulk of head cell populations. Our data accord with previous work; the original LABL study has reported that *per* oscillation in clock neurons dampens with age using the same dvPDF-Gal4 driver^36^. A study using whole-head has shown that the oscillation of clock genes remains robust at 42 days and dampens only at 57 days in an LD cycle^33^, well beyond the 28 day-old flies that we examined on the second day of DD. In contrast, we showed that the FB clock was dampened by both aging and HFD in qRT-PCR and LABL measurements alike. Together, these results identify the FB clock as a principal node at which the effects of aging and HFD converge. In this study, we used male flies to minimize the effects of mating and reproductive status. Because aging and dietary responses are sexually dimorphic^41,42^, further investigation will determine whether the vulnerability of the FB is conserved across sexes.

Both aging and HFD are well-known inducers of chronic inflammation^43,44^, and in *Drosophila* aging is associated with elevated AMP and immune-gene expression^29,45^. The *Drosophila* FB, a principal humoral immune tissue^22^, can be major source of age- and HFD-induced immune activation. Because aging and HFD both dampen the FB clock (Figs. 2, 3), and CLK disruption in the FB upregulated AMP expression (Figs. 5, S2), the dampening of the FB clock by these stressors could contribute to their associated chronic inflammation. Whether the immune activation is harmful has been debated. Previous studies have shown that overexpression of AMPs is cytotoxic and shorten lifespan^46^, whereas AMPs are dispensable in some context^47^. Our data support the previous findings and indicate that the effects of immune activation are conditional, in that CLK disruption shortened lifespan only on HFD (Fig. 4), even though it elevated AMP expression on both SD and HFD (Figs. 5, S2). We therefore propose a model in which immune overactivation induced by CLK dysfunction is necessary but not sufficient to compromise survival, becoming lethal only with the additional metabolic burden of HFD, although the mechanism of this synergy remains to be explored. To establish causality, manipulating AMPs or IMD/Relish signaling in the FB of CLK-disrupted flies with dietary stress will be required to determine whether survival and behavioral phenotypes are restored.

Given that chronic immune activation is associated with behavioral decline across species^39,40^, we asked whether CLK disruption induces behavioral changes. CLK disruption reduced locomotor activity and dampened behavioral rhythmicity (Fig. 6). These data suggest that the behavioral deficits accompany the immune overactivation caused by CLK disruption, although the causal relationship remains to be established. Notably, CLK disruption impaired behavior, similar to aging and HFD (Fig. 1), supporting the idea that aging and HFD converge on the FB clock to impair behavior and physiology. Furthermore, the behavioral changes caused by FB-restricted manipulation of CLK imply non-cell autonomous signaling from the FB to the circuit driving locomotor output. Although how the signals remotely reach the circuit remains to be characterized, the FB is a well-known secretory tissue that delivers cytokines and peptide hormones at a distance^48^. In addition to secreted AMPs, FB-derived peptidergic signals, including Unpaired 2, a secreted peptide functionally analogous to mammalian leptin, can relay nutritional and immune state to the neural circuits controlling behavior^49^. Further investigation is necessary to identify the signals that convey peripheral cues to the brain in order to coordinate behaviors under chronic inflammation.

The most unexpected aspect of our results is that FB-specific disruption of individual core clock components induced different outcomes. Although CLK and CYC act together as the transcriptional activator in the canonical TTFL, and although both CLK and CYC disruption comparably dampened clock gene oscillation in the FB (Fig. S1), only CLK disruption shortened lifespan on HFD, overactivated immunity, and impaired locomotor rhythmicity (Figs. 4–6). If these phenotypes arose simply from the loss of timekeeping function, the manipulations of each clock component should have shown equivalent outcomes. Instead, the differences in their phenotypes suggests that CLK carries an additional function, separable from its canonical timekeeping role, which restrains immune gene expression. In our study, CLK was disrupted with a dominant-negative form of CLK protein that might act beyond simple loss of function. Although the UAS-CLK^DN^ has been widely used and well established^21,50,51^ and the comparison with UAS-CYC^DN^ further strengthens our interpretation, additional experiments will be needed to assess precise molecular dynamics of CLK. Building on emerging evidence^52,53^, mapping CLK’s binding sites across the genome and identifying proteins that interact with CLK at tissue-specific resolution would define how CLK acts in immune and metabolic regulation.

However, non-canonical roles of the core clock components have been increasingly recognized as their non-behavioral roles in peripheral tissues are examined. In addition to its role as a transcriptional activator, mammalian CLOCK acts as a histone acetyltransferase^54^ and regulates NF-kB-dependent transcription independent of its circadian partner BMAL1^55^. BMAL1, in turn, forms a heterodimer with a non-canonical partner, hypoxia-inducible factor 2 alpha, to regulate myocardial injury^56^. In *Drosophila*, CLK, but not PER or TIM, is associated with Relish/ NF-kB signaling in the head^57^. Our findings accord with this trend. As CLOCK acts independently of its canonical partner BMAL1 in mammals, CLK could act independently of CYC in the FB in *Drosophila*. Although our investigation was confined to the FB, future work in other peripheral tissues may reveal a common pattern of non-circadian roles for CLK. Further dissection of the molecular dynamics of each clock component may reveal ways to prevent chronic inflammation that accompanies aging and dietary stress.

Our findings highlight the *Drosophila* FB clock as a principal site at which the detrimental effects of aging and HFD converge, and reveal an additional function of CLK, separable from its CYC-dependent timekeeping role in the TTFL, in restraining inflammation, which is essential for survival on HFD. More broadly, our work offers insight into how the loss of clock robustness in metabolic tissue drives inflammatory disruption that accompanies aging and dietary stress.

## RESOURCE AVAILABILITY

### Lead contact

Requests for further information and resources should be directed to and will be fulfilled by the lead contact, Annika F Barber.

### Material availability

This study did not generate new unique reagents.

### Data and code availability

RNA-seq data have been deposited at GEO (accession number pending). All data to reproduce plots and analysis will be publicly available at Zenodo as of the date of publication.

## ACKNOWLEDGMENTS

We are grateful to Peter Johnstone and Deniz Top for training and assistance in implementing the LABL assay. We thank Cynthia Hsu and Rebecca Moore for assistance with obtaining and documenting fly stocks. We thank all members of the Barber Lab for helpful discussions and comments on the manuscript. We acknowledge the Waksman Institute Busch funds for summer undergraduate support. Stocks obtained from the Bloomington Drosophila Stock Center were used in this study. This study was funded by the NIH R35 GM155111 (to AFB) and American Federation for Aging Research Grants for Junior Faculty 2021 (to AFB).

## AUTHOR CONTRIBUTIONS

Conceptualization, K.F. and A.F.B.; formal analysis, K.F., S.M., V.K., and A.F.B.; funding acquisition, A.F.B.; investigation, K.F., S.M., V.K., C.F., S.S., A.F.B.; supervision, A.F.B.; visualization, K.F. and S.M.; writing – original draft, K.F., S.M., and A.F.B.; writing – review and editing, K.F. and A.F.B.

## METHODS

### Fly food, stocks, and husbandry

Flies were maintained on standard cornmeal medium (BDSC) at 25° C, which was used as the SD in all experiments, under a 12:12 LD cycle. HFD was prepared by adding pork lard to the SD at 20% (w/v)^58^. Briefly, melted pork lard (Epic Provisions) was poured into a graduated cylinder, SD was added to the final volume, and the mixture was homogenized on a magnetic stirrer at 155 °C for 30 minutes, dispensed in 4–5 mL aliquots into vials, and stored at 4° C until use. Unless noted otherwise, flies were transferred to fresh food every 2–3 days. Fly strains are listed in Supplemental file 1. Iso_31_, as isogenic strain of *w*^1118^, was used as a genetic background^59^, and all other strains were back-crossed to Iso_31_ for at least 6 generations.

### DAM assays

Newly eclosed flies (within ∼72 h post-eclosion) were placed in fresh bottles containing SD for 3 days to allow maturation and mating. Male flies were housed at 20 animals per vial until loading. Young (7–9 days post-eclosion) or middle-aged (21–23 days post-eclosion) males were individually placed in glass DAM tubes containing the same food on which they had been maintained and loaded into Drosophila Activity Monitors (TriKinetics, Waltham, MA). For Fig. 1, 5 days in LD or 6 days DD in two independent cohorts were recorded and the first day of DD were excluded from analysis. For Fig. 6, 3 days in LD followed by 6 days in DD were recorded, and the last day of LD and the first day of DD were excluded from analysis. We used ClockLab (Actimetrics, Lafayette, IN) to quantify total daily activity, 24-h behavioral rhythm strength by fast Fourier transform (FFT) and period length by chi-squared periodogram for each fly in constant conditions. Flies not showing rhythmic behavior (FFT < 0.05) were excluded from analysis of period length. We used PHASE to quantify 24 h sleep amount, sleep bout length, and sleep bout number for individual flies in LD^60^.

### Quantitative RT-PCR

Male flies were entrained to 12:12 LD cycle throughout development, transferred to DD on 7–9 or 28–30 days post-eclosion as young and middle-aged flies, then dissected on the second day of DD at 4-h intervals from CT 0 to CT 20. Each biological replicate consisted of 50 heads or 15 FBs from flies maintained on SD or HFD with 3 biological replicates per time point. RNA was extracted using TRIzol™ reagent (Invitrogen, Carlsbad, CA) followed by PureLink RNA Mini Kit (Invitrogen) for head samples or PureLink RNA Micro Kit (Invitrogen) for FB samples, with on-column DNase treatment according to the manufacturer’s protocol. 100 ng of RNA was reverse transcribed with SuperScript IV VILO Master Mix (Invitrogen). qRT-PCR was performed on a CFX Opus 384 Real-Time PCR System (Bio-Rad, Hercules, CA) using iTaq™ Universal SYBR^®^ Green Supermix (Bio-Rad). Cycling conditions were 95° C for 3 m, followed by 40 cycles of 95° C for 10 s and 60° C for 30 s, followed by melt curve analysis. Each sample was run in 3 technical replicates. Relative expression levels were calculated by the ΔΔCt method and normalized to *α-tubulin* (*α-tub*) for head samples and *ribosomal protein L32* (*rpl32*) for FB samples. Rhythmicity and amplitude were assessed using the JTK_Cycle algorithm implemented in the MetaCycle package in R^61^. Primer sequences listed in Supplemental file 2.

### LABL assay

LABL flies, which carry the LABL construct under the control of the minimal *per* promoter, were a gift from Dr. Deniz Top^36^. To express luciferase in specific tissues, LABL flies were crossed with flies carrying *UAS*-FLP2 and a Gal4 driver. *Lsp2*-Gal4 and *dvPDF-*Gal4 were used to target the FB and the LNv and *cryptochrome*^+^ LNd populations of clock neurons respectively. Male flies were housed at a density of 15–20 animals until loading. The day before loading, D-luciferin potassium salt (Research Products International, Mount Prospect, IL) was mixed into SD or HFD to a final concentration of 15 mM. Approximately 1.3 mL of food, a volume sufficient to limit fly movement along the z-axis, was dispensed into custom 35-mm plates (Actimetrics) and stored at 4° C in the dark overnight. Fifteen male flies were loaded into each plate, covered with a 22×30-1 mm coverslip (Fisher Scientific, Pittsburgh, PA), and placed in a LumiCycle 32 color luminometer (Actimetrics). Plates were placed in different positions to ensure equal readout by the four photomultiplier tubes. Luminescence was measured in 6-minute intervals for 9 days in DD using Lumicycle software on a Windows PC. Data was analyzed using custom Python code^36^ (LABLv9.py; https://www.top-lab.org/ downloads or https://github.com/deniztop/LABL). The 9-day recording was used to display overall oscillation, while the first day was excluded with the remaining days were used for period analysis. Amplitude was quantified as the average trough subtracted from the average peak for the first 5 days.

### Survival assay

Newly eclosed males (within ∼72 h post-eclosion) were placed on SD or HFD and monitored throughout their lifespan. Flies were maintained were housed at 15 animals per vial and transferred to fresh vials every 2–3 days, at which point dead flies were recorded.

### RNA-sequencing and analysis

FBs were dissected from *Lsp2*-Gal4>*UAS*-CLK^DN^, *Lsp2*-Gal4>*UAS*-CYC^DN^, and Gal4 parental control (*Lsp2*-Gal4>+) young (7–9 days post-eclosion) males maintained on SD or HFD. Each biological replicate consisted of 20 FBs collected at ZT 4, with 3 replicates per genotype. RNA extraction, library preparation, and sequencing were performed by Admera Health (South Plainfield, NJ). Libraries were sequenced as 150 bp paired-end reads on the Illumina NovaSeq X Plus (10B). Raw reads were subjected to adapter trimming and quality filtering with fastp (v0.23.4)^62^, and read quality was assessed with FastQC (v0.12.1, https://www.bioinformatics.babraham.ac.uk/projects/fastqc/). Filtered reads were mapped to the *Drosophila melanogaster* reference genome (BDGP6) using HISAT2 (v2.2.1)^63^, sorted and indexed with SAMtools (v1.6)^64^. Transcript abundance was counted with StringTie (v2.1.7)^65^ using the Ensemble annotation (release 113; Drosophila_melanogaster.BDGP6.46.113.gtf), which produced gene-level expression in transcripts per million (TPM). The gene-level read count matrix was generated using prepDE.py3. Gene expression was compared, and volcano plots were generated with TCC-GUI^66^. Genes passing a false discovery rate (FDR) < 0.25, log_2_ fold change > 2.5, and an average expression (A value) > 6 were considered as candidate genes. To identify the most relevant hits, we implemented this relaxed FDR combined with stringent cutoffs for fold change and expression abundance. Gene ontology (GO) enrichment analysis was performed with Metascape^67^.

### Statistical analysis

Statistical tests and the value and definition of *n* are reported in the figure legends. Statistical analyses were performed in OriginPro 2025.

## REFERENCES

1. Barzilai, N., Huffman, D.M., Muzumdar, R.H., and Bartke, A. (2012). The Critical Role of Metabolic Pathways in Aging. Diabetes 61, 1315–1322. 10.2337/db11-1300.

2. Baechle, J.J., Chen, N., Makhijani, P., Winer, S., Furman, D., and Winer, D.A. (2023). Chronic inflammation and the hallmarks of aging. Molecular Metabolism 74, 101755. 10.1016/j.molmet.2023.101755.

3. Hood, S., and Amir, S. (2017). The aging clock: circadian rhythms and later life. Journal of Clinical Investigation 127, 437–446. 10.1172/JCI90328.

4. Bass, J., and Takahashi, J.S. (2010). Circadian Integration of Metabolism and Energetics. Science 330, 1349–1354. 10.1126/science.1195027.

5. Scheiermann, C., Kunisaki, Y., and Frenette, P.S. (2013). Circadian control of the immune system. Nat Rev Immunol 13, 190–198. 10.1038/nri3386.

6. Antunes, L.C., Levandovski, R., Dantas, G., Caumo, W., and Hidalgo, M.P. (2010). Obesity and shift work: chronobiological aspects. Nutrition Research Reviews 23, 155–168. 10.1017/S0954422410000016.

7. James, S.M., Honn, K.A., Gaddameedhi, S., and Van Dongen, H.P.A. (2017). Shift Work: Disrupted Circadian Rhythms and Sleep—Implications for Health and Well-being. Curr Sleep Medicine Rep 3, 104–112. 10.1007/s40675-017-0071-6.

8. Leproult, R., Holmbäck, U., and Van Cauter, E. (2014). Circadian Misalignment Augments Markers of Insulin Resistance and Inflammation, Independently of Sleep Loss. Diabetes 63, 1860–1869. 10.2337/db13-1546.

9. Kohsaka, A., Laposky, A.D., Ramsey, K.M., Estrada, C., Joshu, C., Kobayashi, Y., Turek, F.W., and Bass, J. (2007). High-Fat Diet Disrupts Behavioral and Molecular Circadian Rhythms in Mice. Cell Metabolism 6, 414–421. 10.1016/j.cmet.2007.09.006.

10. Kiran, S., Rakib, A., Kodidela, S., Kumar, S., and Singh, U.P. (2022). High-Fat Diet-Induced Dysregulation of Immune Cells Correlates with Macrophage Phenotypes and Chronic Inflammation in Adipose Tissue. Cells 11, 1327. 10.3390/cells11081327.

11. Patke, A., Young, M.W., and Axelrod, S. (2020). Molecular mechanisms and physiological importance of circadian rhythms. Nat Rev Mol Cell Biol 21, 67–84. 10.1038/s41580-019-0179-2.

12. Allada, R., White, N.E., So, W.V., Hall, J.C., and Rosbash, M. (1998). A Mutant *Drosophila* Homolog of Mammalian *Clock* Disrupts Circadian Rhythms and Transcription of *period* and *timeless*. Cell 93, 791–804. 10.1016/S0092-8674(00)81440-3.

13. Rutila, J.E., Suri, V., Le, M., So, W.V., Rosbash, M., and Hall, J.C. (1998). CYCLE Is a Second bHLH-PAS Clock Protein Essential for Circadian Rhythmicity and Transcription of Drosophila period and timeless. Cell 93, 805–814. 10.1016/S0092-8674(00)81441-5.

14. Darlington, T.K., Wager-Smith, K., Ceriani, M.F., Staknis, D., Gekakis, N., Steeves, T.D.L., Weitz, C.J., Takahashi, J.S., and Kay, S.A. (1998). Closing the Circadian Loop: CLOCK-Induced Transcription of Its Own Inhibitors per and tim. Science 280, 1599–1603. 10.1126/science.280.5369.1599.

15. Lee, C., Bae, K., and Edery, I. (1999). PER and TIM Inhibit the DNA Binding Activity of a Drosophila CLOCK-CYC/dBMAL1 Heterodimer without Disrupting Formation of the Heterodimer: a Basis for Circadian Transcription. Molecular and Cellular Biology 19, 5316–5325. 10.1128/MCB.19.8.5316.

16. Crespo-Flores, S.L., and Barber, A.F. (2022). The Drosophila circadian clock circuit is a nonhierarchical network of peptidergic oscillators. Current Opinion in Insect Science 52, 100944. 10.1016/j.cois.2022.100944.

17. Reinhard, N., Fukuda, A., Manoli, G., Derksen, E., Saito, A., Möller, G., Sekiguchi, M., Rieger, D., Helfrich-Förster, C., Yoshii, T., et al. (2024). Synaptic connectome of the Drosophila circadian clock. Nat Commun 15, 10392. 10.1038/s41467-024-54694-0.

18. Yildirim, E., Curtis, R., and Hwangbo, D.-S. (2022). Roles of peripheral clocks: lessons from the fly. FEBS Letters 596, 263–293. 10.1002/1873-3468.14251.

19. Arrese, E.L., and Soulages, J.L. (2010). Insect Fat Body: Energy, Metabolism, and Regulation. Annual Review of Entomology 55, 207–225. 10.1146/annurev-ento-112408-085356.

20. Xu, K., Zheng, X., and Sehgal, A. (2008). Regulation of Feeding and Metabolism by Neuronal and Peripheral Clocks in Drosophila. Cell Metabolism 8, 289–300. 10.1016/j.cmet.2008.09.006.

21. Xu, K., DiAngelo, J.R., Hughes, M.E., Hogenesch, J.B., and Sehgal, A. (2011). The Circadian Clock Interacts with Metabolic Physiology to Influence Reproductive Fitness. Cell Metabolism 13, 639–654. 10.1016/j.cmet.2011.05.001.

22. Lemaitre, B., and Hoffmann, J. (2007). The Host Defense of Drosophila melanogaster. Annual Review of Immunology 25, 697–743. 10.1146/annurev.immunol.25.022106.141615.

23. Wang, C., Lutes, L.K., Barnoud, C., and Scheiermann, C. (2022). The circadian immune system. Science Immunology 7, eabm2465. 10.1126/sciimmunol.abm2465.

24. Lee, J.-E., and Edery, I. (2008). Circadian Regulation in the Ability of *Drosophila* to Combat Pathogenic Infections. Current Biology 18, 195–199. 10.1016/j.cub.2007.12.054.

25. Stone, E.F., Fulton, B.O., Ayres, J.S., Pham, L.N., Ziauddin, J., and Shirasu-Hiza, M.M. (2012). The Circadian Clock Protein Timeless Regulates Phagocytosis of Bacteria in Drosophila. PLOS Pathogens 8, e1002445. 10.1371/journal.ppat.1002445.

26. Trindade de Paula, M., Poetini Silva, M.R., Machado Araujo, S., Cardoso Bortolotto, V., Barreto Meichtry, L., Zemolin, A.P.P., Wallau, G.L., Jesse, C.R., Franco, J.L., Posser, T., et al. (2016). High-Fat Diet Induces Oxidative Stress and MPK2 and HSP83 Gene Expression in Drosophila melanogaster. Oxidative Medicine and Cellular Longevity 2016, 4018157. 10.1155/2016/4018157.

27. Eickelberg, V., Rimbach, G., Seidler, Y., Hasler, M., Staats, S., and Lüersen, K. (2022). Fat Quality Impacts the Effect of a High-Fat Diet on the Fatty Acid Profile, Life History Traits and Gene Expression in Drosophila melanogaster. Cells 11, 4043. 10.3390/cells11244043.

28. Stobdan, T., Sahoo, D., Azad, P., Hartley, I., Heinrichsen, E., Zhou, D., and Haddad, G.G. (2019). High fat diet induces sex-specific differential gene expression in Drosophila melanogaster. PLOS ONE 14, e0213474. 10.1371/journal.pone.0213474.

29. Zerofsky, M., Harel, E., Silverman, N., and Tatar, M. (2005). Aging of the innate immune response in Drosophila melanogaster. Aging Cell 4, 103–108. 10.1111/j.1474-9728.2005.00147.x.

30. Rhodenizer, D., Martin, I., Bhandari, P., Pletcher, S.D., and Grotewiel, M. (2008). Genetic and environmental factors impact age-related impairment of negative geotaxis in *Drosophila* by altering age-dependent climbing speed. Experimental Gerontology 43, 739–748. 10.1016/j.exger.2008.04.011.

31. Wang, R., Yin, Y., Li, J., Wang, H., Lv, W., Gao, Y., Wang, T., Zhong, Y., Zhou, Z., Cai, Y., et al. (2022). Global stable-isotope tracing metabolomics reveals system-wide metabolic alternations in aging Drosophila. Nat Commun 13, 3518. 10.1038/s41467-022-31268-6.

32. Koh, K., Evans, J.M., Hendricks, J.C., and Sehgal, A. (2006). A *Drosophila* model for age-associated changes in sleep:wake cycles. Proc. Natl. Acad. Sci. U.S.A. 103, 13843–13847. 10.1073/pnas.0605903103.

33. Luo, W., Chen, W.-F., Yue, Z., Chen, D., Sowcik, M., Sehgal, A., and Zheng, X. (2012). Old flies have a robust central oscillator but weaker behavioral rhythms that can be improved by genetic and environmental manipulations. Aging Cell 11, 428–438. 10.1111/j.1474-9726.2012.00800.x.

34. Liao, S., Amcoff, M., and Nässel, D.R. (2021). Impact of high-fat diet on lifespan, metabolism, fecundity and behavioral senescence in Drosophila. Insect Biochemistry and Molecular Biology 133, 103495. 10.1016/j.ibmb.2020.103495.

35. Nayak, N., and Mishra, M. (2021). High fat diet induced abnormalities in metabolism, growth, behavior, and circadian clock in *Drosophila melanogaster*. Life Sciences 281, 119758. 10.1016/j.lfs.2021.119758.

36. Johnstone, P.S., Ogueta, M., Akay, O., Top, I., Syed, S., Stanewsky, R., and Top, D. (2022). Real time, in vivo measurement of neuronal and peripheral clocks in Drosophila melanogaster. eLife 11, e77029. 10.7554/eLife.77029.

37. Maurange, C., and Lanet, E. (2014). Building a brain under nutritional restriction: insights on sparing and plasticity from Drosophila studies. Front. Physiol. 5. 10.3389/fphys.2014.00117.

38. Poe, A.R., Xu, Y., Zhang, C., Lei, J., Li, K., Labib, D., and Han, C. (2020). Low FoxO expression in Drosophila somatosensory neurons protects dendrite growth under nutrient restriction. eLife 9, e53351. 10.7554/eLife.53351.

39. Biesmans, S., Meert, T.F., Bouwknecht, J.A., Acton, P.D., Davoodi, N., Haes, P.D., Kuijlaars, J., Langlois, X., Matthews, L.J.R., Donck, L.V., et al. (2013). Systemic Immune Activation Leads to Neuroinflammation and Sickness Behavior in Mice. Mediators of Inflammation 2013, 271359. 10.1155/2013/271359.

40. Arora, S., Critchley, G., Dekmak, A.S., Miesenböck, G., Kempf, A., and Ligoxygakis, P. (2025). Loss of the NF-κB negative regulator Pirk in Drosophila links brain and gut immunity to neurodegeneration. Brain Commun 7, fcaf144. 10.1093/braincomms/fcaf144.

41. Regan, J.C., Khericha, M., Dobson, A.J., Bolukbasi, E., Rattanavirotkul, N., and Partridge, L. (2016). Sex difference in pathology of the ageing gut mediates the greater response of female lifespan to dietary restriction. eLife 5, e10956. 10.7554/eLife.10956.

42. Wu, Q., Yu, G., Cheng, X., Gao, Y., Fan, X., Yang, D., Xie, M., Wang, T., Piper, M.D.W., and Yang, M. (2020). Sexual dimorphism in the nutritional requirement for adult lifespan in Drosophila melanogaster. Aging Cell 19, e13120. 10.1111/acel.13120.

43. Hotamisligil, G.S. (2006). Inflammation and metabolic disorders. Nature 444, 860–867. 10.1038/nature05485.

44. Franceschi, C., Garagnani, P., Parini, P., Giuliani, C., and Santoro, A. (2018). Inflammaging: a new immune–metabolic viewpoint for age-related diseases. Nat Rev Endocrinol 14, 576–590. 10.1038/s41574-018-0059-4.

45. Pletcher, S.D., Macdonald, S.J., Marguerie, R., Certa, U., Stearns, S.C., Goldstein, D.B., and Partridge, L. (2002). Genome-Wide Transcript Profiles in Aging and Calorically Restricted *Drosophila melanogaster*. Current Biology 12, 712–723. 10.1016/S0960-9822(02)00808-4.

46. Badinloo, M., Nguyen, E., Suh, W., Alzahrani, F., Castellanos, J., Klichko, V.I., Orr, W.C., and Radyuk, S.N. (2018). Overexpression of antimicrobial peptides contributes to aging through cytotoxic effects in Drosophila tissues. Archives of Insect Biochemistry and Physiology 98, e21464. 10.1002/arch.21464.

47. Hanson, M.A., and Lemaitre, B. (2023). Antimicrobial peptides do not directly contribute to aging in Drosophila, but improve lifespan by preventing dysbiosis. Dis Model Mech 16, dmm049965. 10.1242/dmm.049965.

48. Meschi, E., and Delanoue, R. (2021). Adipokine and fat body in flies: Connecting organs. Molecular and Cellular Endocrinology 533, 111339. 10.1016/j.mce.2021.111339.

49. Rajan, A., and Perrimon, N. (2012). *Drosophila* Cytokine Unpaired 2 Regulates Physiological Homeostasis by Remotely Controlling Insulin Secretion. Cell 151, 123–137. 10.1016/j.cell.2012.08.019.

50. Tanoue, S., Krishnan, P., Krishnan, B., Dryer, S.E., and Hardin, P.E. (2004). Circadian Clocks in Antennal Neurons Are Necessary and Sufficient for Olfaction Rhythms in Drosophila. Current Biology 14, 638–649. 10.1016/j.cub.2004.04.009.

51. Hodge, B.A., Meyerhof, G.T., Katewa, S.D., Lian, T., Lau, C., Bar, S., Leung, N.Y., Li, M., Li-Kroeger, D., Melov, S., et al. (2022). Dietary restriction and the transcription factor clock delay eye aging to extend lifespan in Drosophila Melanogaster. Nat Commun 13, 3156. 10.1038/s41467-022-30975-4.

52. Meireles-Filho, A.C.A., Bardet, A.F., Yáñez-Cuna, J.O., Stampfel, G., and Stark, A. (2014). *cis*-Regulatory Requirements for Tissue-Specific Programs of the Circadian Clock. Current Biology 24, 1–10. 10.1016/j.cub.2013.11.017.

53. Mahesh, G., Rivas, G.B.S., Caster, C., Ost, E.B., Amunugama, R., Jones, R., Allen, D.L., and Hardin, P.E. (2020). Proteomic analysis of Drosophila CLOCK complexes identifies rhythmic interactions with SAGA and Tip60 complex component NIPPED-A. Sci Rep 10, 17951. 10.1038/s41598-020-75009-5.

54. Doi, M., Hirayama, J., and Sassone-Corsi, P. (2006). Circadian Regulator CLOCK Is a Histone Acetyltransferase. Cell 125, 497–508. 10.1016/j.cell.2006.03.033.

55. Spengler, M.L., Kuropatwinski, K.K., Comas, M., Gasparian, A.V., Fedtsova, N., Gleiberman, A.S., Gitlin, I.I., Artemicheva, N.M., Deluca, K.A., Gudkov, A.V., et al. (2012). Core circadian protein CLOCK is a positive regulator of NF-κB–mediated transcription. Proceedings of the National Academy of Sciences 109, E2457–E2465. 10.1073/pnas.1206274109.

56. Ruan, W., Li, T., Bang, I.H., Lee, J., Deng, W., Ma, X., Luo, C., Du, F., Yoo, S.-H., Kim, B., et al. (2025). BMAL1–HIF2A heterodimer modulates circadian variations of myocardial injury. Nature 641, 1017–1028. 10.1038/s41586-025-08898-z.

57. Liu, X., Fang, J., Cheng, D., Luan, W., Lv, Y., Hu, W., Pan, L., and Zhang, Y. The Circadian Transcription Factor CLOCK Modulates Oxidative Stress Resistance via the ACHL–Relish Axis in Drosophila. Advanced Science n/a, e14388. 10.1002/advs.202514388.

58. Havula, E., Ghazanfar, S., Lamichane, N., Francis, D., Hasygar, K., Liu, Y., Alton, L.A., Johnstone, J., Needham, E.J., Pulpitel, T., et al. (2022). Genetic variation of macronutrient tolerance in Drosophila melanogaster. Nat Commun 13, 1637. 10.1038/s41467-022-29183-x.

59. Ryder, E., Blows, F., Ashburner, M., Bautista-Llacer, R., Coulson, D., Drummond, J., Webster, J., Gubb, D., Gunton, N., Johnson, G., et al. (2004). The DrosDel Collection: A Set of P-Element Insertions for Generating Custom Chromosomal Aberrations in Drosophila melanogaster. Genetics 167, 797–813. 10.1534/genetics.104.026658.

60. Persons, J.L., Abhilash, L., Lopatkin, A.J., Roelofs, A., Bell, E.V., Fernandez, M.P., and Shafer, O.T. (2022). PHASE: An Open-Source Program for the Analysis of DrosophilaPhase, Activity, and Sleep Under Entrainment. J Biol Rhythms 37, 455–467. 10.1177/07487304221093114.

61. Wu, G., Anafi, R.C., Hughes, M.E., Kornacker, K., and Hogenesch, J.B. (2016). MetaCycle: an integrated R package to evaluate periodicity in large scale data. Bioinformatics 32, 3351–3353. 10.1093/bioinformatics/btw405.

62. Chen, S., Zhou, Y., Chen, Y., and Gu, J. (2018). fastp: an ultra-fast all-in-one FASTQ preprocessor. Bioinformatics 34, i884–i890. 10.1093/bioinformatics/bty560.

63. Kim, D., Paggi, J.M., Park, C., Bennett, C., and Salzberg, S.L. (2019). Graph-based genome alignment and genotyping with HISAT2 and HISAT-genotype. Nat Biotechnol 37, 907–915. 10.1038/s41587-019-0201-4.

64. Danecek, P., Bonfield, J.K., Liddle, J., Marshall, J., Ohan, V., Pollard, M.O., Whitwham, A., Keane, T., McCarthy, S.A., Davies, R.M., et al. (2021). Twelve years of SAMtools and BCFtools. Gigascience 10, giab008. 10.1093/gigascience/giab008.

65. Pertea, M., Pertea, G.M., Antonescu, C.M., Chang, T.-C., Mendell, J.T., and Salzberg, S.L. (2015). StringTie enables improved reconstruction of a transcriptome from RNA-seq reads. Nat Biotechnol 33, 290–295. 10.1038/nbt.3122.

66. Su, W., Sun, J., Shimizu, K., and Kadota, K. (2019). TCC-GUI: a Shiny-based application for differential expression analysis of RNA-Seq count data. BMC Res Notes 12, 133. 10.1186/s13104-019-4179-2.

67. Zhou, Y., Zhou, B., Pache, L., Chang, M., Khodabakhshi, A.H., Tanaseichuk, O., Benner, C., and Chanda, S.K. (2019). Metascape provides a biologist-oriented resource for the analysis of systems-level datasets. Nat Commun 10, 1523. 10.1038/s41467-019-09234-6.

