## Supplemental Material for "Fat body CLOCK restrains innate immunity and maintains survival under dietary stress in *Drosophila*"

### Figure S1

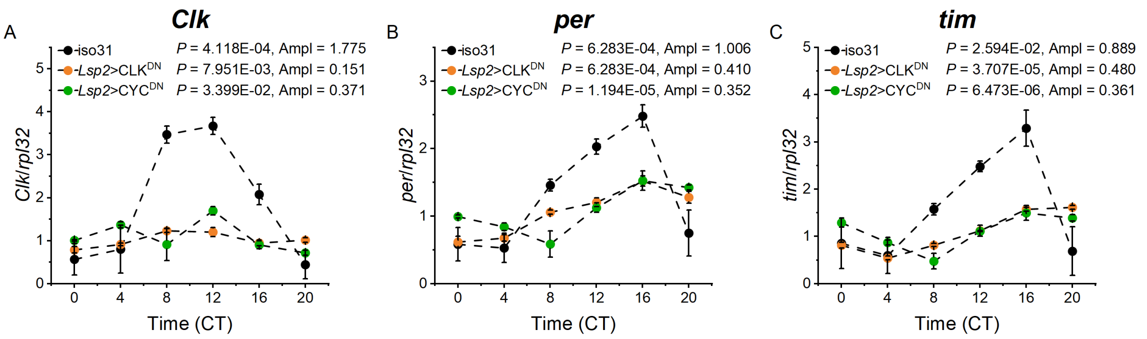


**Fig. S1 Rhythmic transcription of core clock genes is dampened in the fat body of CLK$^{DN}$ or CYC$^{DN}$ flies.**

Daily oscillation of clock gene transcripts in the fat body (FB; A–C) of young (7–9 days old) male iso$^{31}$ (control; black), *Lsp2*-Gal4>*UAS*-CLK$^{DN}$ (orange), and *Lsp2*-Gal4>*UAS*-CYC$^{DN}$ (green) flies maintained on a standard diet. The iso31 data are replotted from Fig. 2 D–F for comparison. mRNA expression of *Clock* (C*lk*; A), *period* (*per*; B), and *timeless* (*tim*; C) was measured by qRT-PCR across circadian time with clock mutants showing reduced gene expression oscillation amplitude compared to the isogenic control. Expression in the FB was normalized to *ribosomal protein L32* (*rpl32)*. Data represent the mean ± SEM of 3 biological replicates per time point; each FB replicate comprised 15 FBs. Rhythmicity (*P*) and amplitude (Ampl) were determined by JTK_Cycle.

### Figure S2

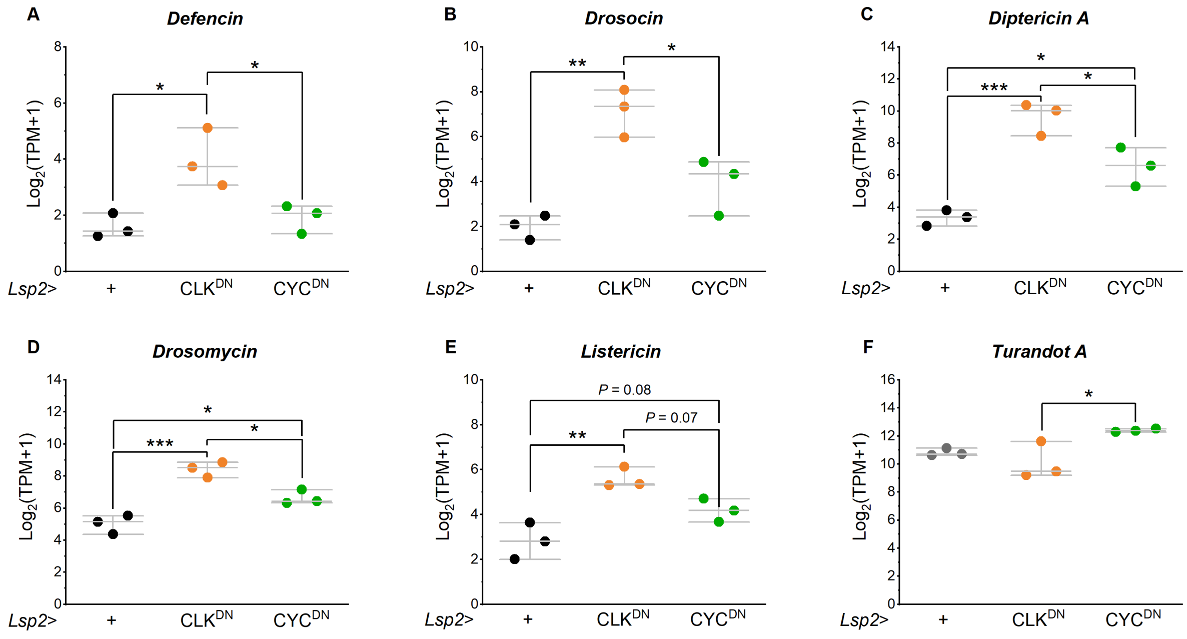


**Fig. S2 Disruption of CLOCK upregulates AMP transcription.** Expression of antimicrobial peptides and immune genes: *Defensin* (A), *Drosocin* (B), *Diptericin* A (C), *Drosomycin* (D), *Listericin* (E), and *Turandot A* (F) in the fat body of young (7–9 days old) male *Lsp2*-Gal4 controls (black), *Lsp2*-Gal4>*UAS*-CLK$^{DN}$ (orange), and *Lsp2*-Gal4>*UAS*-CYC$^{DN}$ (green) flies maintained on a standard diet. All immune genes were elevated in *Lsp2*-Gal4>*UAS*-CLK$^{DN}$ except *Turandot A*, which was reduced, whereas *Lsp2*-Gal4>*UAS*-CYC$^{DN}$ had no significant effect. Top and bottom horizontal lines indicate interquartile range (25$^{th}$–75$^{th}$ percentiles), the central line is the median, and each point is one biological replicate. $n$ = 3 per genotype; each FB replicate comprised 20 FBs. Data were analyzed by one-way ANOVA followed by Tukey's *post hoc* test. *, $P < 0.05$; **, $P < 0.01$; ***, $P < 0.001$

### Supplemental file 1

**Supplemental file 1.** Genotypes of flies used in the figures. Iso$_{31}$ strain of $w^{1118}$ was used as a genetic background and all other strains were backcrossed to Iso$_{31}$ for at least 6 generations.

| Figure | Label | Genotype |
|---|---|---|
| Figure 1 | - | *w*; +; + |
| Figure 2 | - | *w*; +; + |
| Figure 3A–H | - | *w*; LABL/+; *Lsp2*-Gal4/*UAS*-FLP2 |
| Figure 3I–P | - | *w*; *dvPDF*-Gal4/+; LABL/*UAS*-FLP2 |
| Figure 4A, E | *Lsp2*>CLK$^{DN}$ | *w*; *UAS*-CLK$^{DN}$/+; *Lsp2*-Gal4/+ |
| | *Lsp2*>+ | *w*; +/+; *Lsp2*-Gal4/+ |
| | +>CLK$^{DN}$ | *w*; *UAS*-CLK$^{DN}$/+; +/+ |
| Figure 4B, F | *Lsp2*>CYC$^{DN}$ | *w*; *UAS*-CYC$^{DN}$/+; *Lsp2*-Gal4/+ |
| | *Lsp2*>+ | *w*; +/+; *Lsp2*-Gal4/+ |
| | +>CYC$^{DN}$ | *w*; *UAS*-CYC$^{DN}$/+; +/+ |
| Figure 4C, G | *Lsp2*>*per*$^{CRISPR}$ | *w*; *UAS*-*per*-gRNA/+; *Lsp2*-Gal4/*UAS*-Cas9 |
| | *Lsp2*>+ | *w*; +/+; *Lsp2*-Gal4/+ |
| | +>*per*$^{CRISPR}$ | *w*; *UAS*-*per*-gRNA/+; *UAS*-Cas9/+ |
| Figure 4D, H | *Lsp2*>*tim*$^{CRISPR}$ | *w*; *UAS*-*tim*-gRNA/+; *Lsp2*-Gal4/*UAS*-Cas9 |
| | *Lsp2*>+ | *w*; +/+; *Lsp2*-Gal4/+ |
| | +>*tim*$^{CRISPR}$ | *w*; *UAS*-*tim*-gRNA/+; *UAS*-Cas9/+ |
| Figure 5A, B, D–J | *Lsp2*>CLK$^{DN}$ | *w*; *UAS*-CLK$^{DN}$/+; *Lsp2*-Gal4/+ |
| | *Lsp2*>CYC$^{DN}$ | *w*; *UAS*-CYC$^{DN}$/+; *Lsp2*-Gal4/+ |
| | *Lsp2*>+ | *w*; +/+; *Lsp2*-Gal4/+ |
| Figure 5C, D | *Lsp2*>CLK$^{DN}$ | *w*; *UAS*-CLK$^{DN}$/+; *Lsp2*-Gal4/+ |
| | *Lsp2*>+ | *w*; +/+; *Lsp2*-Gal4/+ |
| Figure 6 | *Lsp2*>CLK$^{DN}$ | *w*; *UAS*-CLK$^{DN}$/+; *Lsp2*-Gal4/+ |
| | *Lsp2*>CYC$^{DN}$ | *w*; *UAS*-CYC$^{DN}$/+; *Lsp2*-Gal4/+ |
| | *Lsp2*>+ | *w*; +/+; *Lsp2*-Gal4/+ |
| | +>CLK$^{DN}$ | *w*; *UAS*-CLK$^{DN}$/+; +/+ |
| | +>CYC$^{DN}$ | *w*; *UAS*-CYC$^{DN}$/+; +/+ |
| Figure S1 | Iso$_{31}$ | Iso$_{31}$ |
| | *Lsp2*>CLK$^{DN}$ | *w*; *UAS*-CLK$^{DN}$/+; *Lsp2*-Gal4/+ |
| | *Lsp2*>CYC$^{DN}$ | *w*; *UAS*-CYC$^{DN}$/+; *Lsp2*-Gal4/+ |

### Supplemental file 1

| | | |
|---|---|---|
| Figure S2 | *Lsp2*>CLK$^{DN}$<br>*Lsp2*>CYC$^{DN}$<br>*Lsp2*>+ | *w*; *UAS*-CLK$^{DN}$/+; *Lsp2*-Gal4/+<br>*w*; *UAS*-CYC$^{DN}$/+; *Lsp2*-Gal4/+<br>*w*; +/+; *Lsp2*-Gal4/+ |

### Supplemental file 2

**Supplemental file 2. Primer sequences used for qRT-PCR**

| **Gene** | | **Sequence** |
|---|---|---|
| *$\alpha$-tub* | Fw | CGTCTGGACCACAAGTTCGA |
| | Rv | CCTCCATACCCTCACCAACGT |
| *rpL32* | Fw | GCCCAAGGGTATCGACAACA |
| | Rv | GCGCTTGTTCGATCCGTAAC |
| *Clk* (for the head) | Fw | TTCTCGATGGTGTTCTCGGTG |
| | Rv | AGTTCGCAAAGCCAACGG |
| *Clk* (for the FB) | Fw | GGATAAGTCCACGGTCCTGA |
| | Rv | CTCCAGCATGAGGTGAGTGT |
| *per* (for the head) | Fw | CCTGAAAGACGCGAT GGTG |
| | Rv | CGTCAATCCATGGTCCCG |
| *per* (for the FB) | Fw | TCCCAATCCGCGTACAACAA |
| | Rv | GGCACCTTCTTCGTCATGGA |
| *tim* (for the head and FB) | Fw | AGGAGGATGAAGATGAGGACGAAGT |
| | Rv | GCCCAATTTGACCGCAGATAC |
